# Schizophrenia and human self-domestication: an evolutionary linguistics approach

**DOI:** 10.1101/072751

**Authors:** Antonio Benítez-Burraco, Lorena Di Pietro, Marta Barba, Wanda Lattanzi

## Abstract

Schizophrenia (SZ) is a pervasive neurodevelopmental disorder entailing social and cognitive deficits, including marked problems with language. Its complex multifactorial etiopathogenesis, including genetic and environmental factors, is still widely uncertain. SZ incidence has always been high and quite stable in human populations, across time and regardless of cultural implications, due to unclear reasons. It has been hypothesised that SZ pathophysiology may involve the biological components that changed during the recent human evolutionary history and led to our distinctive mode of cognition, which includes language skills. In this paper we explore this possibility, focusing on the self-domestication of the human species. This has been claimed to account for many human-specific distinctive traits, including aspects of our behaviour and cognition, and to favour the emergence of complex languages through cultural evolution. The “domestication syndrome” in mammals comprises the constellation of traits exhibited by domesticated strains, seemingly resulting from the hypofunction of the neural crest. It is our intention to show that people with SZ exhibit more marked domesticated traits at the morphological, physiological, and behavioural levels. We also show that genes involved in domestication and neural crest development and function comprise nearly 20% SZ candidates, most of which exhibit altered expression profiles in the brain of SZ patients, specifically in areas involved in language processing. Based on these observations, we conclude that SZ may represent an abnormal ontogenetic itinerary for the human faculty of language, resulting, at least in part, from changes in genes important for the “domestication syndrome” and, primarily involving the neural crest.

## 1. Introduction

Schizophrenia (SZ) is a pervasive neurodevelopmental condition entailing different and severe social and cognitive deficits. Core distinctive symptoms of SZ include delusions, hallucinations, impaired motivation, reduction in spontaneous speech, and social withdrawal; cognitive impairment, episodes of elated mood, and episodes of depressive mood are also commonly observed (van Os and Kapur 2009, Owen et al., 2016). SZ prevalence has been found stable across time and cultures, to the extent that it has been considered a human-specific disease. Indeed, susceptibility genes are poorly conserved across species, some of them being absent in great apes (Brüne, 2004, Pearlson and Folley, 2008). Also, most of the biological components that seem to have played a central role in the evolution of human cognition are found impaired in SZ patients. For instance, after our split from great apes, the frontal cortical circuitry was remodelled; this circuitry is responsible for many human-specific cognitive abilities and is found dysfunctional in patients with SZ and other psychiatric conditions (Teffer and Semendeferi, 2012). Likewise, genomic regions that have undergone positive selection in anatomically-modern humans (AMHs) are enriched in gene loci associated with SZ (Srinivasan et al., 2016). This enrichment has been recently linked to functional elements like introns and untranslated regions (Srinivasan et al., 2017). This rises the intriguing possibility that most of SZ risk alleles appeared more recently in human evolution. Overall, these evidences suggest that the evolutionary changes occurred in the human lineage, particularly after the split from extinct hominins, may help clarifying some aspects of SZ. On the other hand, delving into the SZ polygenic etiopathogenesis, which acts synergistically with unclear environmental factors, might help understand the changes that brought about our human distinctive cognitive phenotype, including our language abilities.

Language deficits are a hallmark of SZ, which has been defined as “the price that homo sapiens pays for language” (Crow, 2000). These usually manifest as problems in speech perception (in the form of auditory verbal hallucinations), abnormal speech production (known as Formal Thought Disorder, FTD), and production of abnormal linguistic content (that is, delusions) (Stephane et al., 2007, and 2014). These major positive symptoms can be reduced to disturbances in linguistic computation (Hinzen and Roselló 2015) that result from atypical brain development and wiring during growth (Li et al., 2009, and 2012). In our previous work we have showed that this abnormal mode of processing language can be specifically drawn back to an abnormal, distinctive oscillatory profile of the brain during language computation (Murphy and Benítez-Burraco, 2016a). Also, we have showed that candidates for SZ are overrepresented among the genes believed to be involved in the evolution of our language-readiness, that is, our species-specific ability to learn and use languages (Boeckx and Benítez-Burraco, 2014a, 2014b; Benítez-Burraco and Boeckx, 2015, Murphy and Benítez-Burraco, 2016a).

Besides the genomic and epigenomic changes that favoured our speciation, we expect that our cognitive phenotype was also modelled by changes occurred later, during our self-domestication (Benítez-Burraco et al., 2016a). The idea of human beings as domesticated primates can be drawn back to Darwin (1871). Recent comparisons with extinct hominins have revealed that AMHs exhibit a number of domesticated traits, including differences in the brain and the face, changes in dentition, reduction of aggressiveness, and retention of juvenile characteristics (see Thomas, 2014 for details). Many authors have argued that the relaxation of the selective pressures on our species resulting from this process of self-domestication may have contributed to the creation of the cultural niche that favoured the emergence of modern languages (Hare and Tomasello, 2005, Deacon, 2009, Thomas, 2014, among others). This niche provides humans with an extended socialization window, enabling them to receive a greater amount of linguistic stimuli, to involve in enhanced and prolonged communication exchanges with other conspecifics, and to experiment with language for a longer time. In particular, language complexity is expected to increase in these comfortable conditions, as attested by domestic strains of songbirds, in which domestication triggers variation and complexity in their songs (Takahasi and Okanoya, 2010, Kagawa et al., 2012). Importantly, this possibility is supported by several linguistic studies revealing positive correlations between aspects of linguistic complexity and aspects of social complexity (Wray and Grace, 2007, Lupyan and Dale, 2010), or pointing out to the emergent nature of core properties of human languages, resulting from cultural transmission (Benítez-Burraco, 2016).

Several selectionist accounts of why humans became self-domesticated have been posited over time, ranging from selection against aggression and towards social tolerance, to a by-product of mate-choices, to adaptation to the human-made environment (Thomas, 2014). In our recent work we have hypothesised that self-domestication might be (also) a by-product of the changes that brought about our more globular skull/brain and our language-readiness (Benítez-Burraco et al., 2016a). The reason is that candidates for globularization and language-readiness are found among (and interact with) the genes believed important for the development and function of the neural crest (NC). And as noted by Wilkins et al (2014), the set of traits observed in domestic mammals, ranging from changes in the craniofacial region, to changes in the skin, the reproductive and vital cycles, and behaviour (the so-called ‘domestication syndrome”), may result from the hypofunction of the NC, in turn triggered by the selection for tameness (see Sánchez-Villagra et al., 2016, for a recent account).

Building on this hypothesis, in our previous work we have showed that the complex pathophysiology of some human cognitive diseases entailing problems with language can be, at least in part, linked to an abnormal presentation of the “domestication syndrome” (Murphy and Benítez-Burraco, 2016a). Specifically, we have discussed how patients suffering from autism spectrum disorders (ASD) exhibit a plethora of distinctive behavioural, neurological, and physical anomalies, including dysmorphic features, that seem to be opposite as the “domesticated” traits observed in typically-developing (TD) individuals (Benítez-Burraco et al., 2016b).

Interestingly, ASD and SZ have been hypothesised to be opposite poles in the continuum of cognitive modes, encompassing also the TD one. Their opposed natures can be tracked from brain structure and function to neurodevelopmental paths, to cognitive abilities (Crespi and Badcock, 2008). We have previously shown as well that SZ and ASD patients process language differently, and exhibit distinctive, disorder-specific oscillatory profiles when computing language (Murphy and Benítez-Burraco, 2016b). Similarly to SZ, language deficits in ASD can be linked to many of the changes occurred during our speciation, to the extent that candidates for this condition are also overrepresented among the genes believed to account for the evolution of our language-readiness (see Benítez-Burraco and Murphy, 2016).

In this paper we wish to explore the possibility that SZ patients exhibit exacerbated, disease-specific signatures of the “domestication syndrome”. If we are right, our hypothesis could pave the way towards exploring the etiopathogenesis of SZ, and related language impairment, under an original standpoint. To this aim, we begin providing a general account of the domesticated traits found in SZ patients. Thereafter, we will focus on the molecular etiopathogenesis of SZ and check whether genes that have been found involved in the domestication process are somehow represented among SZ candidates. Considering the relevant role of the neural NC in the domestication process, we also consider, in this search, genes involved in NC development and function. Through this comparative evaluation of candidates, we will define a subset of overlapping genes involved in both SZ and in domestication and/or NC. The functional role of these selected genes will be discussed in detail, focusing on their contribution to etiopathogenesis of SZ and their implication for cognition and language. In addition, in order to evaluate their actual functional involvement, we will delve into their differential expression profiles in the SZ brain, by *in silico* analysis of previously published data.

Environmental factors are known to contribute to SZ too (see Brown, 2011; Geoffroy et al., 2013; Moran et al., 2016, for reviews). Likewise, although domestication has genetic roots (as originally suggested by Wilkins et al. and as we will show here), domestication results as well in the creation of a cultural niche that favours the maintenance of domestic features by cultural evolution. Accordingly, although our focus is put on the genes, the role of the environment cannot be dismissed. The same is true for language indeed, as suggested above. Modern language seemingly evolved as the result of changes in brain genes that brought about a differential cognitive ability (aka *language-readiness*) and favoured a domesticated phenotype in our species, but it is also a consequence of subsequent changes in the (proto)linguistic systems, that were triggered and facilitated by the domestic environment in which human beings are reared.

## 2. Domestication features in SZ

Most of the features observed in the “domestication syndrome” described by Wilkins and colleagues (2014) are found generally exacerbated in SZ individuals (Figure 1). Domestic varieties of mammals exhibit a distinctive set of common traits (Figure 1, top), including neoteny, shorter reproductive cycles, depigmentation, and increased tameness. Changes in behaviour seemingly result from reduced levels of stress hormones (including adrenocorticoids, adrenocorticotropic hormone, cortisol, and corticosterone), and particularly, from the delay in the maturation of the adrenal glands, which also gives rise to an increase of the duration of the immaturity of the hypothalamic-pituitary-adrenal system (the HPA axis) and a hypofunction of the sympathetic nervous system, which provides the animal with a longer socialization window. Many of the differences with their wild conspecifics concern to the craniofacial area. These include changes in ear size and shape, changes in the orofacial area (including shorter snouts and smaller jaws), changes in dentition (particularly, smaller teeth), and a reduced brain capacity (specifically, of components of the forebrain such as the amygdala or parts of the limbic system) (see Wilkins et al., 2014 and Sánchez-Villagra et al., 2016 for details). Below we provide with a detailed description of domestication traits commonly found in schizophrenic patients (Figure 1, down).

**Figure 1.**
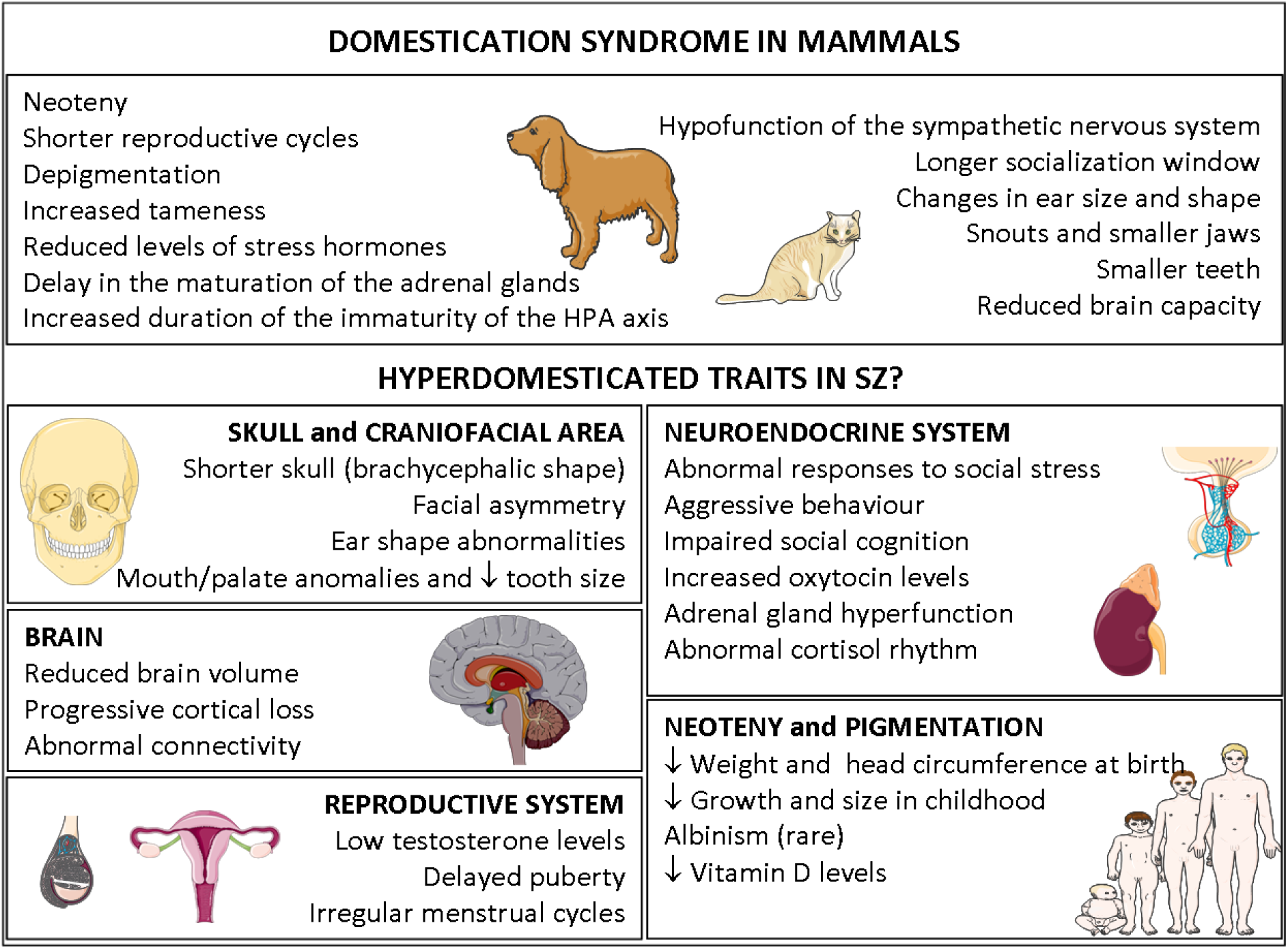
Schizophrenia and the domestication syndrome. The diagram is meant to symbolize the anomalous presentation of the “domestication syndrome” in people with schizophrenia. Main features observed in domesticated mammals (Wilkins et al., 2014, Sánchez-Villlagra, 2016) are shown in the upper box, while selected clinical findings from the SZ spectrum, that may resemble «hyper-domesticated traits», are categorized below. Pictures were gathered and modified from “Slide kit Servier Medical Art” (available at www.servier.com).

### Physical anomalies

Minor dysmorphisms are typically featured in the craniofacial area of SZ patients. Indeed, facial asymmetries, particularly those arising along the midfacial junctions (between frontonasal and maxillary prominence derivatives), are reproducibly found in these patients (Gourion et al., 2004, Deutsch et al., 2015). Additionally, ear shape abnormalities (including adherent ear lobes, lower edges of the ears extending backward/upward, malformed ears, asymmetrical ears, or cuspidal ears) are usually observed in SZ phenotypes (Yoshitsugu et al., 2006, Akabaliev et al., 2011, Lin et al., 2012). Some of these features (like prominent crux of helix and ear lobe crease, or primitive shape of the ear) are considered as pathognomonic for SZ in the differential diagnosis of psychotic conditions (Trixler et al., 2001, Praharaj et al., 2012). Anomalies in the mouth (e.g. decreased tooth size, abnormal palate shape and size) are also commonly observed in schizophrenics (Ismail et al., 1998, Rajchgot et al., 2009, Hajnal et al., 2016). Likewise, the odds of having a psychotic disorder seem to be increased in people with shorter and wider palates (McGrath et al., 2002). More generally, the odds of having a psychotic disorder seem to be increased in people with smaller lower-facial heights (glabella to subnasal) (McGrath et al., 2002). Some studies suggest a significant association between minor physical anomalies and the early onset of the disease (Hata et al., 2003).

### Brain anomalies and dysfunctions

Metanalyses of structural neuroimaging studies in the SZ brain are indicative of a significant reduction of total brain volume, which mostly affects to the hippocampus, the thalamus and the cortex, most pronounced in the frontal and temporal lobes (Steen et al., 2006, Haijma et al., 2013, Haukvik et al., 2013). This is seemingly due to the impairment of the surface expansion of the cortex during brain growth, which impacts more on the left hemisphere and results in a relative areal contraction of diverse functional networks (Palaniyappan et al., 2011). Gray matter reduction in SZ is also associated with longer duration of illness and reduced sensitivity to antipsychotic medications. Accordingly, brain volume constraints in SZ are better explained as a combination of early neurodevelopmental disturbance and disease progression (Haijma et al., 2013). Metanalyses of longitudinal neuroimaging studies of the schizophrenic brain further suggest that SZ entails a disorder-specific trajectory of morphological change (compared to other similar conditions like bipolar disorder), which is characterised by a progressive grey matter loss confined to fronto-temporal cortical regions (De Peri et al., 2012, Liberg et al., 2016). Children with childhood onset SZ revels that SZ is characterised by reduced cerebral volume and cortical thickness during childhood and adolescence, which is levelled off in adulthood, as well as by deficits in local connectivity and increased long-range connectivity (Baribeau and Anagnostou, 2013).

The reduction of brain volume is expected to impact cognitive and language abilities of patients and to account for distinctive symptoms of the disease (see also section 5). Specifically, schizophrenic patients with FTD show clusters of volume reduction in the medial frontal and orbitofrontal cortex bilaterally (related to poverty of content of speech), and in two left-sided areas approximating to Broca’s and Wernicke’s areas (related to the fluent disorganization component of FTD) (Sans-Sansa et al., 2013). Likewise, reduced brain activity in the left pars triangularis of Broca’s area positively correlates with volume reduction of this area (Iwashiro et al., 2016). Interestingly, antipsychotic-naive patients show more pronounced volume reductions in caudate nucleus and thalamus (Haijma et al., 2013), which play a key role in language processing (Murphy, 2015). Finally, we wish highlight that amygdala volume is usually reduced in schizophrenics (Li et al., 2015, Okada et al., 2016, Rich et al., 2016).

### Behavioural traits and neuroendocrine impairment

Aggressive behaviour, being involved in the behavioural traits of the “domestication syndrome”, is frequent in SZ, and paranoid belief may associate with it (Darrell-Berry et al., 2016). Interestingly, no positive correlation seems to exist between physical aggression and neuropsychological performance in patients (unless they have attained severe impairment that induces constant uncontrollable outbursts) (Lapierre et al., 1995).

SZ involves as well an impairment of social cognition. Oxytocin is a neuropeptide hormone that, within a wide range of organic functions, is able to affect social interactions and response to social stimuli at various levels (reviewed by Romano et al., 2016). Specifically, it has been recently argued to modulate the multimodality that characterizes our higher-order linguistic abilities (Theofanopoulou, 2016). Oxytocin promotes social play in domestic dogs and the appropriate use of human social cues (Oliva et al., 2015, Romero et al., 2015). A positive correlation between the SZ progression and oxytocin levels in the central nervous system has been observed (Beckmann et al., 1985), which is plausibly explained by a decreased sensitivity to the hormone (Strauss et al., 2015, Glovinsky et al., 1994, Sasayama et al., 2012). Treatment with oxytocin indeed improves verbal memory learning tasks in SZ patients (Feifel et al., 2012), and attenuates the negative symptoms of the disease (Feifel et al., 2010, Modabbernia et al., 2013, Gibson et al., 2014, Davis et al., 2014).

The hypothalamus-pituitary axis (HPA) is also affected in SZ, with both hyper- and hypofunction being described (Bradley and Dinan, 2010). Accordingly, heightened cortisol levels are observed in patients with SZ, especially in those who are not medicated (Walker et al., 2008). At the same time, Hempel et al., (2010) found that cortisol concentration in the plasma decreases more markedly during the day in SZ patients than in healthy controls, and that the decrease of HPA axis sensitivity correlates with the severity of negative symptoms. In male patients, diagnosed with first-episode SZ, higher afternoon cortisol levels at the beginning of medical treatment are related to impaired memory performance (Havelka et al., 2016). Girshkin et al., (2016) found that SZ patients do not show significant differences in waking cortisol levels, in the cortisol awakening response, or in immediate post-cortisol awakening control decline compared to controls. However, they also found that they exhibit a significant absence of the increase in cortisol responsivity to stress. According to Ciufolini et al., (2014), SZ is characterised by an attenuated HPA axis response to social stress: despite a normal cortisol production rate, schizophrenics have lower cortisol levels than controls, both in anticipation and after exposure to social stress. In the TD population, HPA activity increases around puberty, with a postpubertal rise in baseline cortisol secretion linked with pubertal stage (Walker et al., 2001, Gunnar et al., 2009). It has been suggested that delayed adrenarche correlates with a higher risk for SZ (Saugstad 1989a, 1989b).

### Other features

With regard to neoteny, it is noteworthy that SZ patients exhibit lower weight and reduced head circumference at birth (Cannon et al., 2002), along with slower growth rates and smaller sizes in childhood (Gunnell et al., 2003, Haukka et al., 2008).

Reproductive cycles are also affected in both male and female SZ patients. Delayed age at puberty is associated with greater severity of negative SZ prodromal symptoms in males (Ramanathan et al., 2015). In women higher negative symptom scores and greater functional impairment correlate with later age of menarche (Hochman and Levine 2004). Nearly 50% of women with SZ have irregular menses that are frequently associated to low levels of oestradiol, although no differences in their neuropsychological status have been found compared to patients with regular menses (Gleeson et al., 2016). There is ample evidence of the protective effect of estradiol with respect to SZ, because it interacts with the neurotransmitter systems implicated in the disease, and because it enhances cognition and memory, and reverses the symptoms (Gogos et al., 2015). Men with SZ have, indeed, lower levels of testosterone than healthy controls, and an inverse correlation between serum testosterone and negative symptoms of the disease has been described (Ramsey et al., 2013, Sisek-Šprem et al., 2015). However, in more aggressive patients this correlation is not found (Sisek-Šprem et al., 2015). Interestingly, circulating testosterone levels in schizophrenic males predict performance on verbal memory, processing speed, and working memory (Moore et al., 2013). Men with SZ show a less pronounced activation of the middle frontal gyrus when inhibiting response to negative stimuli, and this response is inversely related to testosterone level, contrary to what is observed in healthy subjects (Vercammen et al., 2013). Testosterone significantly affects brain development, particularly targeting the hypothalamus, the amygdala, and the hippocampus, and impacting on aspects of memory consolidation (Filová et al., 2013).

Lastly, concerning changes in pigmentation, an association between SZ and albinism has been occasionally reported (Clarke and Buckley, 1989). In turn, hyperpigmentation is typically described as a side effect of neuroleptic drugs (specifically, of phenothiazines) used in SZ treatment (Otreba et al., 2015). Interestingly, low serum vitamin D levels have been found in SZ patients and they correlate with the severity of psychotic symptoms (Yüksel et al., 2014). The molecular background for this link may rely on shared features of latitude-adaptation observed in both SZ- and vitamin D-related genes, which suggest that SZ etiopathogenesis may encounter latitude dependent adaptive changes in vitamin D metabolism (Amato et al., 2010).

As noted in section 1, the constellation of symptoms that characterize the “domestication syndrome” have been hypothesised to result as the unselected by-product of a reduce input in NC cells (Wilkins et al., 2014). The phenotypical presentation of human neurocristopathies commonly includes features that have been described in domesticated mammals (Sánchez-Villagra et al., 2016). Interestingly, well-defined neurocristopathies like velocardiofacial syndrome (OMIM#192430) and Di George syndrome (OMIM#188400), involve schizophrenic features (Mølsted et al., 2010, Zhang et al., 2014, Escot et al., 2016). Likewise, given the NC derivation of most craniofacial structures, craniofacial abnormalities observed in SZ are believed to result from disturbances in the neuroectoderm development, hence representing putative external biomarkers of atypical brain growth (Comptom et al., 2007, Aksoy-Poyraz et al., 2011), and suggesting an additional connection between SZ and domestication, at the level of NC functional implication.

## 3. Genetic signature of domestication/neural crest features in the SZ molecular background.

In order to delve into the molecular background of our hypothesis, we first assessed whether genes that are involved in SZ etiopathogenesis are represented among candidates for domestication and NC development and function. To this aim, we gathered an extended an up-to-date list of SZ-associated genes, through literature mining and database search (using the Schizophrenia Database, http://www.szdb.org/). The list includes 2689 genes with different levels of evidence: genes bearing pathogenic SNPs, genes found mutated in familial forms of the disease, genes resulting from candidate gene approaches and functional studies, genes resulting from GWA and CNV/exome sequencing studies, and genes showing alternative methylation patterns (see the entire list and corresponding details in Supplemental file 1). Regarding candidates for domestication, we have implemented an enlarged list of candidates, which includes the core set of genes proposed by Wilkins and colleagues (2014), along with additional candidates derived from genetic studies performed in different species. The entire list of domestication candidates comprises 127 genes, detailed in Supplemental file 2. Finally, the third gene list considered in the analysis includes genes related to NC development and function. This list comprises 89 genes (see Supplemental file 3) gathered using functional and pathogenic criteria: NC markers, neurochristopathy-associated genes annotated in the OMIM database, genes that are functionally involved in NC induction and specification, genes involved in NC signalling (within NC-derived structures), and genes involved in cranial NC differentiation.

To search the intersections among the three different sets of genes we have employed a simple Venn diagram (Figure 2), drawn using the software designed and made available as a webtool, by the bioinformatics evolutionary genomics group, at the University of Gent (Belgium; webpage: bioinformatics.psb.ugent.be/webtools/Venn/). A Venn diagram shows all possible logical relations between a finite collection of different sets; the diagram consists of multiple overlapping circles, each representing a set.

**Figure 2.**
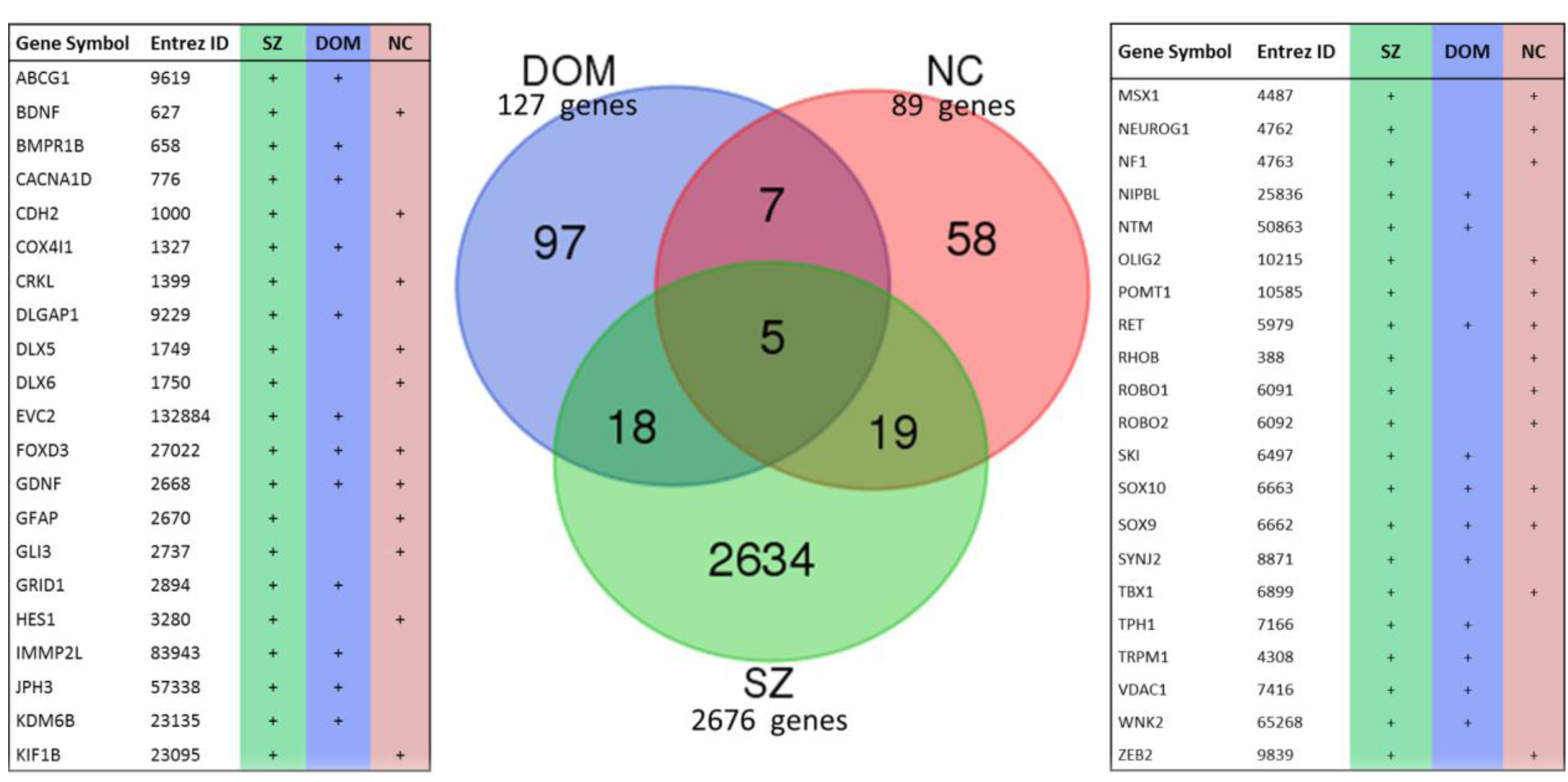
Genetic overlap among schizophrenia, domestication, and neural crest signatures. Venn diagrams show the intersection among the extended list of genes considered as either genomic or functional candidates for schizophrenia (SZ, see Supplemental file 1), domestication (DOM, see Supplemental file 2) and neural crest (NC, see Supplemental file 3). See text for details.

As shown in Figure 2, an overall number of 42 genes were found among the intersection sets of the diagram. In particular, 5 genes (*FOXD3, RET, SOX9, SOX10* and *GDNF*) were overlapping within the three gene lists, hence could represent a selected core of candidates that support our hypothesis. In addition, 18 genes were shared between the SZ list and the domestication list, while 19 genes were shared between SZ and NC candidates. Overall, we found out that over 18% (23 out of 127) of domestication candidates, and 27% (24 out of 89) of genes involved in NC development and function, are listed within those that have been documented as playing a role, as either putative candidates for, or functionally related to SZ. Considering domestication (n: 127) and neural crest (n: 89) candidates altogether, this list comprises 19.4% (42 of 216) SZ candidates.

Below we provide with a brief functional characterization and biological interpretation of these genes.

## 4. Functional implication and biological interpretation of domestication/NC genes in the SZ brain.

We expected that the 42 genes we highlight here as part of the shared signature of domestication and/or NC and SZ (see Figure 2) are functionally interconnected and map on to specific signaling cascades, regulatory pathways, or aspects of brain development and function, of interest for SZ etiopathogenesis, and specifically, for language deficits in this condition. Accordingly, we have employed the license free software for gene network analysis String 10 (www.string-db.org). This allowed predicting quite robust links among the genes. In particular, the core genes (*FOXD3, GDNF, RET, SOX9*, and *SOX10*) are all reciprocally interconnected, hence displayed in the central part of the network (Figure 3). These genes are indeed involved in different steps of NC development and neural specification.

**Figure 3.**
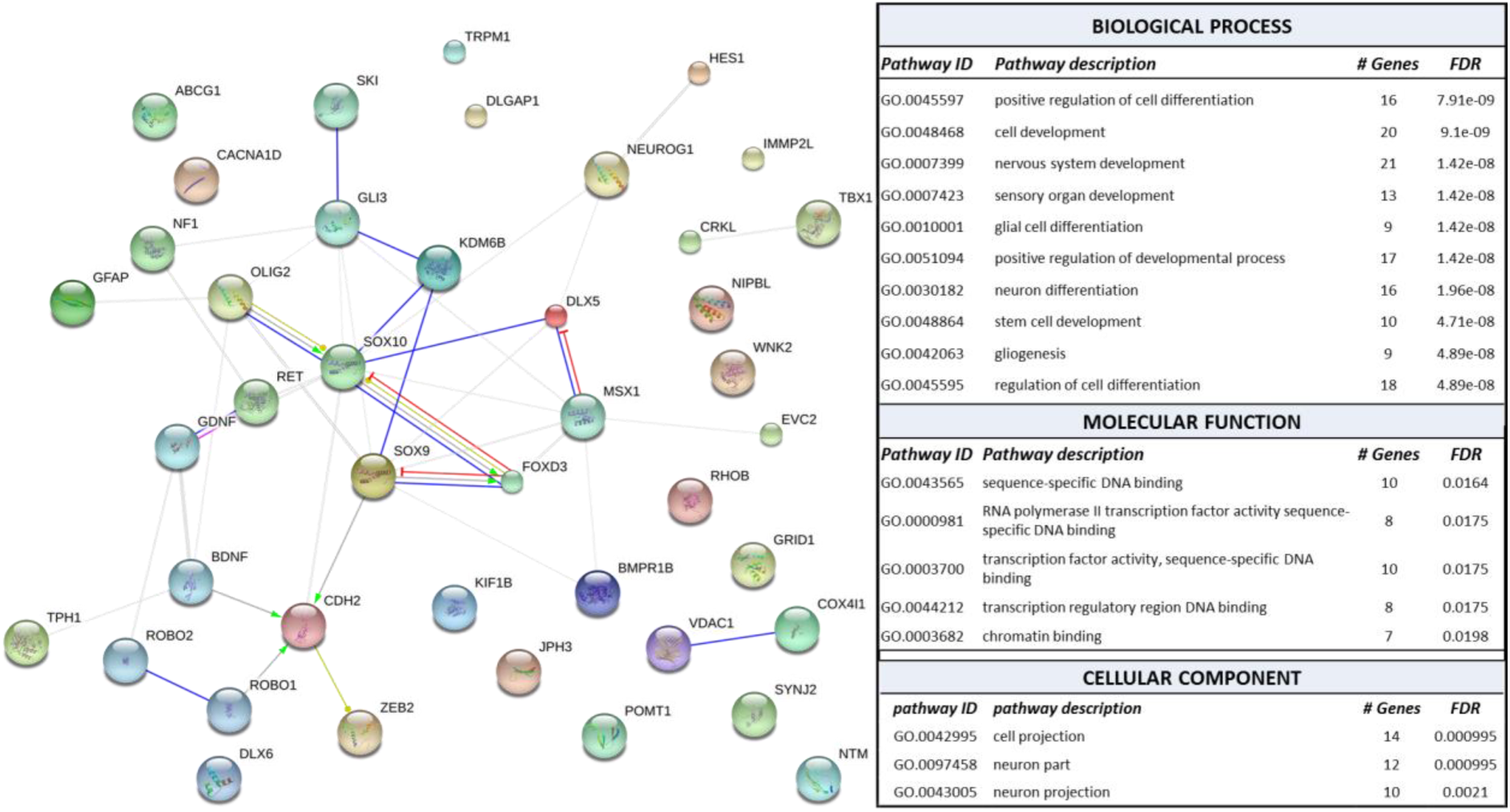
Gene interaction network. The diagram shows the network of known and predicted interactions among genes proposed as candidates for SZ and domestication and/or NC development and function (genes with positive tags in the last two right-sided columns in Figure 2). The network was drawn with String (version 10.0; Szklarczyk et al., 2015)) license-free software (http://string-db.org/), using the molecular action visualization. Colored nodes symbolize gene/proteins included in the query (small nodes are for proteins with unknown 3D structure, while large nodes are for those with known structures). The color of the edges represent different kind of known protein-protein associations. Green: activation, red: inhibition, dark blue: binding, light blue: phenotype, dark purple: catalysis, light purple: post-translational modification, black: reaction, yellow: transcriptional regulation. Edges ending in an arrow symbolize positive effects, edges ending in a bar symbolize negative effects, whereas edges ending in a circle symbolize unspecified effects. Grey edges symbolize predicted links based on literature search (co-mentioned in PubMed abstracts). Stronger associations between proteins are represented by thicker lines. The medium confidence value was .0400 (a 40% probability that a predicted link exists between two enzymes in the same metabolic map in the KEGG database: http://www.genome.jp/kegg/pathway.html). The diagram only represents the potential connectivity between the involved proteins, which has to be mapped onto particular biochemical networks, signaling pathways, cellular properties, aspects of neuronal function, or cell-types of interest. Functional enrichment of the entire gene set, according to Gene Ontology (GO) consortium annotations, was performed using String algorithm for gene network analysis; the output is provided in the table on the right. FDR: false-discovery rate, obtained after Bonferroni correction. A FDR cutoff of 0.05 was set to select significant functions. For the “biological process” and “molecular function” annotations, only the top ten scoring categories are displayed.

*RET* gene is a core domestication candidate (Wilkins et al., 2014), which encodes a tyrosine-protein kinase involved in NC development, and is found deleted in SZ patients (Glessner et al., 2010). *SOX* genes encode master transcriptional regulators of cell-fate programming during development. In particular, *SOX9* and *SOX10* are members of the soxE group, involved in NC development and differentiation (Cheung and Briscoe, 2003). *SOX9* acts specifically in craniofacial development, downstream of WNT and BMP pathways (Liu et al., 2013b), and has been found reproducibly upregulated in SZ brains (Shao and Vawter, 2008, Chen et al., 2013, Lanz et al., 2014). *SOX10* regulates NC stem cells balance during development; it is functionally related to the SZ-susceptibility gene *DISC1* during early NC cell migration (Drerup et al., 2009), and oligodendrocyte differentiation (Hattori et al., 2014). *SOX10* is found hypermethylated in the brain of SZ patients (Iwamoto et al., 2005) and carries SNPs that have been related to the age of onset of clinical manifestations (Yuan et al., 2013). *RET* and *SOX10* are co-expressed as reliable markers in enteric nervous system, although their functional interaction is not clear (Hetz et al., 2014). In turn, *FOXD3* is required during NC development and regulates dorsal mesoderm development in the zebrafish (Chang and Kessler, 2010). The gene locus maps within one of the AMH-specific differentially-methylated genomic regions (Gokhman et al., 2014), suggesting that changes in the gene functional status may have occurred during recent human evolution. *FOXD3* is also a transcriptional target of the SZ-candidate *DISC1*, and is involved in cranial NC migration and differentiation (Drerup et al., 2009). *SOX9* and *SOX10* indirectly interact through *FOXD3*, as shown in Figure 3, being coexpressed at different stages of NC development. In particular, *SOX9* seems to drive the expression of both *SOX10* and *FOXD3* (Cheung and Briscoe, 2003). Finally, *GDNF* is also a core candidate for the domesticated phenotype in mammals (Wilkins et al., 2014). It encodes a neurospecific factor involved in the differentiation of dopaminergic neurons and in synaptogenesis (Christophersen et al., 2007, Ledda et al., 2007). GDNF levels are lower in SZ patients compared with healthy controls (Tunca et al., 2015). With regard to gene networking, GDNF contributes to the activation of *RET* protein-tyrosine kinase (Jing et al., 1996), mediating neuronal survival (Coulpier et al., 2002).

Other genes in the network are directly or indirectly related to the core genes described above. In particular, *MSX1* and *DLX5* homeotic genes also appear to play pivotal roles in this gene network, as they establish functional interactions with many of the candidates shown in Figure 3. Interestingly, *MSX* genes control the spatial organization of the NC-derived craniofacial skeleton (Attanasio et al., 2013, Khadka et al., 2006, Han et al., 2007, Gitton et al., 2011). Also *DLX* homeobox genes function in early NC development, and also in late specification of NC-derived structures (McLarren et al., 2003, Ruest et al., 2003), and play key roles as well in skull and brain development (Jones and Rubinstein, 2004, Kraus and Lufkin, 2006, Vincentz et al., 2016). In particular, *MSX1* encodes a transcriptional repressor specifically involved in odontogenesis (Alappat et al., 2003, Cohen, 2000), hence mutated in orofacial clefting and tooth agenesis (Liang et al., 2016). *MSX1* is a direct downstream target of *DLX5* during early inner ear development (Sajan et al., 2011). Methylation changes in *MSX1* are found in the hippocampus of SZ patients, as a part of the circuit-specific DNA methylation changes affecting the glutamate decarboxylase 1 regulatory network in SZ, which may explain GABAergic dysfunctions in this condition (Ruzicka et al., 2015).

Among the other shared candidates between SZ and domestication, *KDM6B* and *BMPR1B* are shown to interact with *SOX9* and *SOX10* (see Figure 3). *KDM6B* encodes a histone demethylase, which plays a central role in regulation of posterior development. In particular, it activates neuronal gene expression during postnatal and adult brain neurogenesis in the subventricular zone (Park et al., 2014). A frameshift mutation, with unknown functional consequences, has been found in this gene, through whole exome sequencing (WES) of SZ family trios (Fromer et al., 2014). In turn, *BMPR1B* encodes a receptor for bone morphogenetic proteins (BMPs), that are pleiotropic morphogens acting in both bone and neural development. The interaction with *SOX9* is well characterized during postnatal chondrogenesis (Jing et al., 2014). Type Ib BMP receptors also mediate the rate of commissural axon extension in the developing spinal cord (Yamauchi et al., 2013), and, in mice, are involved in supraxial nervous functions (Caronia et al., 2010). *BMPRIB* is annotated among the putative SZ epigenetic signature genes, resulting from genome-wide methylome studies (Aberg et al., 2014).

Also *GLI3* is widely functionally involved in this network. This gene encodes a key mediator of the hedgehog signalling in vertebrates, acting as a repressor in dorsal brain regions (Haddad-Tóvolli et al., 2012). It controls cortical size by regulating the primary cilium-dependent neuronal migration (Wilson et al., 2012). With regards to SZ, a *de novo* missense mutation in *GLI3* has been recently identified through WES (Fromer et al., 2014). *GLI3* has been found to interact, both *in vivo* and *in vitro*, with *SKI* (Ravasi et al., 2010). This gene encodes a transcription factor that regulates TGFβ signaling and is mutated in nearly 90% of cases of Shprintzen-Goldberg syndrome (OMIM#182212), a condition entailing craniofacial and brain anomalies (Au et al., 2014). This gene is found methylated in SZ patients (Montano et al., 2016). Indeed, in the brain, *SKI* regulates the proliferation and differentiation of neural precursors, along with the specification of cortical projections (Lyons et al., 1994, Baranek and Atanasoski 2012). In the mouse embryo, *Ski* is expressed in the migrating NCCs and in NC derivatives (Lyons et al., 1994).

*OLIG2* encodes a transcription factor that interacts with SOX proteins, and plays a key role in the regulation of ventral neuroectodermal progenitor cell fate. Specifically, it is essential for oligodendrocyte function, whose impairment is thought to be a primary pathogenic event in SZ, specifically affecting the prefrontal cortex (Georgieva et al., 2006, Mauney et al., 2015). Indeed, in our network *OLIG2* is also linked to *GFAP*, which encodes a hallmark structural component of mature astrocytes. Interestingly, *GFAP* levels are upregulated in the left posterior superior temporal gyrus (Wernicke’s area) of schizophrenics (Martins de Souza et al., 2009).

Another key node in the network is represented by the *CDH2* gene, which encodes a member of the cadherin superfamily involved in the formation of cartilage and bone, the establishment of left-right asymmetry, and the development of the nervous system (Kadowaki et al., 2007, Martínez-Garay et al., 2016)). Inactivation of *CDH2* in the dorsal telencephalon results in a “double cortex” phenotype, with heterotopic gray matter interposed between zones of white matter (Gil-Sanz et al., 2014). *CDH2* indeed regulates the proliferation and differentiation of ventral midbrain dopaminergic progenitors, the organization of excitatory and inhibitory synaptic circuits, and long-term potentiation in the adult hippocampus (Bozdagi et al., 2010, Sakane and Miyamoto 2013, Nikitczuk et al., 2014). The gene was found mutated in schizophrenics in a WES study (Purcell et al., 2016). *CDH2* integrates *SOX9* signaling and regulates *OLIG2* in neuroepithelial lineage cells, during vertebrate brain development (Sasai et al., 2014). Also, CDH2 cooperates with the BDNF in the aggregation, assembly and mobilization of synaptic vesicles (Bury and Sabo, 2014). *BDNF* encodes a nerve growth factor needed for neuronal survival and synaptic plasticity. Common variants/polymorphisms of the gene have been associated with specific cognitive processes (see Goldberg and Weinberger 2004, González-Giraldo et al., 2014, Jasińska et al., 2016, Wegman et al., 2016) and with the cognitive performance of people suffering from neuropsychiatric conditions. In particular, abnormally low levels of BDNF have been detected in schizophrenics (Palomino et al., 2006). Functional genomics has indeed identified *BDNF* among a list of reliable SZ candidates, contributing to the genetic background for the neurodevelopmental abnormalities leading to the disrupted connectivity occurring in the disease (Ayalew et al., 2012). Being implicated in axonogenesis and neural cell polarization, *CDH2* is also indirectly related to *ROBO* genes. *ROBO1* and *ROBO2* encode highly conserved transmembrane receptors that function in axon guidance and neuronal precursor cell migration. Mutations in *ROBO* genes have been linked to human neurodevelopmental disorders, as discussed in the next section. Noticeably, functional genomics analyses have identified both *ROBO1* and *ROBO2* as candidate loci for SZ risk (Potkin et al., 2009, 2010). Finally, *ZEB2* appears as an additional functional partner of *CDH2* in the network (Figure 3). This gene encodes a transcriptional factor, which acts as an essential regulator of neuroectoderm and NC development, contributing to the development of the neocortex and the hippocampus (Hegarty et al., 2015). Besides being mutated in the Mowat-Wilson syndrome (OMIM#235730), a condition characterized by Hirschsprung disease, craniofacial dismorphisms, and intellectual disability (Adam et al., 2006, Garavelli et al., 2016), *ZEB2* maps within a SZ-associated locus, thus possibly representing a player in the polygenic etiology of this condition (Ripke et al., 2013, Bigdeli et al., 2016).

Other candidate genes are not clearly functionally interconnected in the core interacting network (see Figure 3), although most of them play relevant roles in cognitive functions that have evolved in humans. Most of them are further discussed in section 5 because of their impact on language development.

The functional enrichment, based on gene ontology (GO) annotations, of the gene list considered in this study (Figure 2), point out that most of these genes act in signaling pathways known to be impaired in SZ and might play biological functions that are affected in this condition (see GO annotation table in Figure 3). Noticeably, the top-scoring functional categories, resulting from the functional annotations, include regulation of nervous system development and of cell differentiation, specifically of glial cells and neurons. Among the molecular function GO categories, transcription regulation hits as the most relevant; indeed, many of the genes listed here encode transcription factors and epigenetic modulators, that on their turn modulate the expression of genes with pleiotropic role in brain development, cognitive abilities (including language processing). Finally, considering the cellular localization of the proteins, most of them appear to localize inside the cell projection components, confirming their role as regulators of neuron interconnection.

We have further attempted to delve into the actual functional implication of these selected genes in the SZ molecular pathogenesis, by assessing whether their expression is significantly modulated in the SZ brain. To this aim, we surveyed the Gene Expression Omnibus (GEO) repository (https://www.ncbi.nlm.nih.gov/gds) searching for gene expression datasets obtained from SZ brain profiling. The following dataset were selected and corresponding data were gathered: GSE53987 (prefrontal cortex, hippocampus, and associative striatum; Lanz et al, 2014), GSE4036 (cerebellum; Perrone-Bizzozzero, unpublished data), GSE21935 (temporal cortex; Barnes et al, 2011); GSE35977 (parietal cortex; Chen et al, 2013); GSE62191 (frontal cortex; de Baumont et al, 2015). The specified datasets were selected based on their homogeneous and comparable study design. Additional details are provided in Table 1. Specifically, all datasets were obtained by means of genome-wide microarray expression profiling of dissected cadaveric brain tissues from SZ patients and matched controls. Raw datasets were individually analyzed *in silico* as previously described (Benítez-Burraco et al., 2016). Briefly, the GEO2R tool (http://www.ncbi.nlm.nih.gov/geo/geo2r/; Barrett et al., 2013) was used to compare patients-versus-controls normalized probeset intensities provided by the submitters. The p-value was adjusted, whenever appropriate, using Bonferroni-Hochberg correction for false discovery rate (FDR). A p-value cutoff <0.05 was set for filtering data. Base 2 logarithm transformation of fold changes (logFC) were applied to obtain relative expression changes between SZ patients and corresponding controls.

**Table 1.**
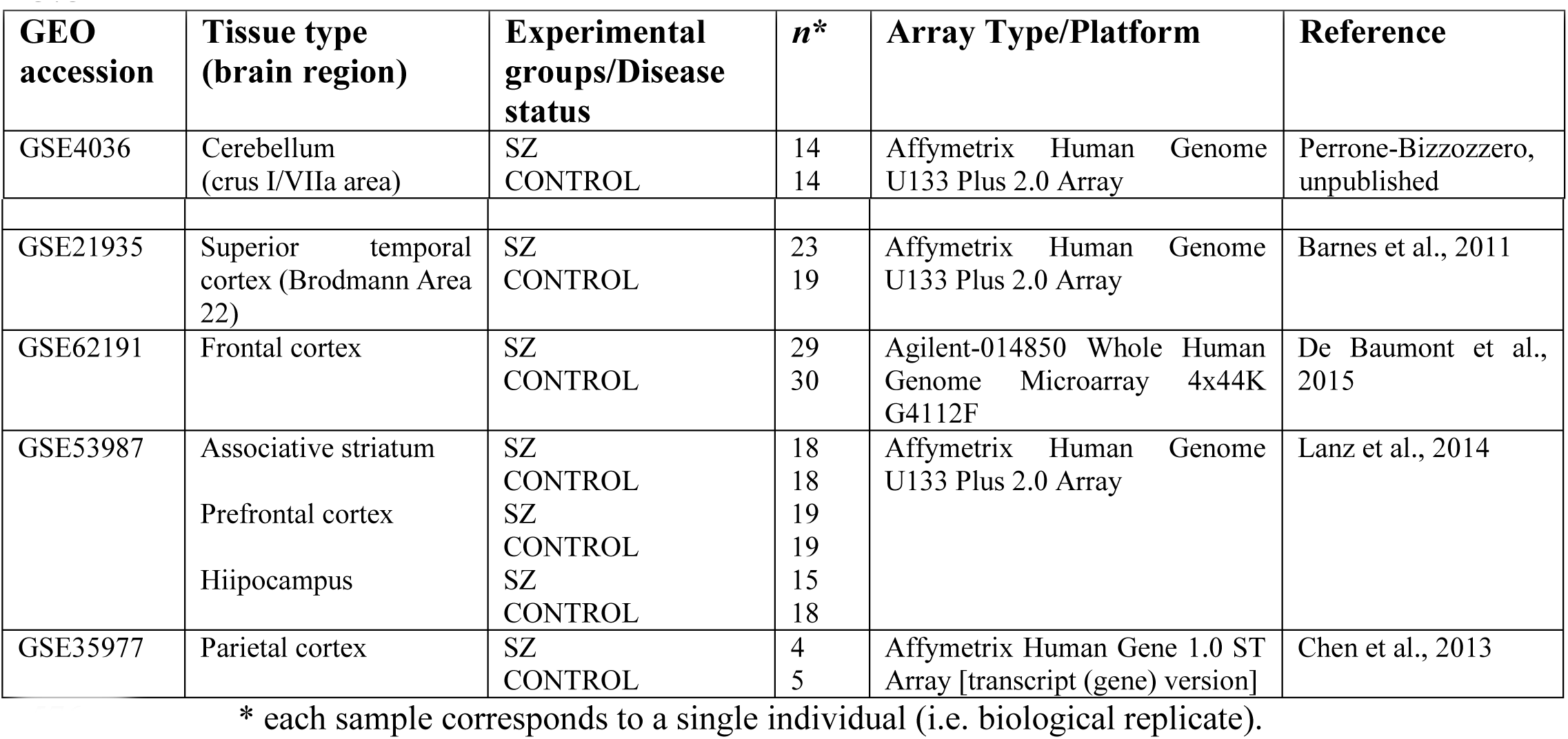
Dataset description.

| GEO accession | Tissue type (brain region) | Experimental groups/Disease status | n* | Array Type/Platform | Reference |
| --- | --- | --- | --- | --- | --- |
| GSE4036 | Cerebellum (crus I/VIIa area) | SZ<br>CONTROL | 14<br>14 | Affymetrix Human Genome U133 Plus 2.0 Array | Perrone-Bizzozzero, unpublished |
| GSE21935 | Superior temporal cortex (Brodmann Area 22) | SZ<br>CONTROL | 23<br>19 | Affymetrix Human Genome U133 Plus 2.0 Array | Barnes et al., 2011 |
| GSE62191 | Frontal cortex | SZ<br>CONTROL | 29<br>30 | Agilent-014850 Whole Human Genome Microarray 4x44K G4112F | De Baumont et al., 2015 |
| GSE53987 | Associative striatum<br>Prefrontal cortex<br>Hippocampus | SZ<br>CONTROL<br>SZ<br>CONTROL<br>SZ<br>CONTROL | 18<br>18<br>19<br>19<br>15<br>18 | Affymetrix Human Genome U133 Plus 2.0 Array | Lanz et al., 2014 |
| GSE35977 | Parietal cortex | SZ<br>CONTROL | 4<br>5 | Affymetrix Human Gene 1.0 ST Array [transcript (gene) version] | Chen et al., 2013 |

\* each sample corresponds to a single individual (i.e. biological replicate).

Through this approach, we have obtained cortical area-specific expression profiles of the SZ brain. Complete gene lists of differentially expressed genes obtained from each dataset are provided in Supplemental file 4. Briefly, the cerebellum expression profiles of SZ patients included in the GSE4036 data series yielded over 1800 annotated genes showing statistically significant (p-value<0.05) differential expression over controls. The data analysis of the temporal cortex data included in the GSE21935 dataset yielded nearly 1950 annotated genes (p<0.05). In the frontal cortex (GSE62191 dataset), the expression profile included 727 differentially expressed annotated genes (p<0.05), whereas in the prefrontal cortex, analyzed in an independent study (GSE53987 dataset), it included nearly 6200 genes (p<0.05). Finally, the in silico analysis of data included in the GSE53987 dataset allowed also obtaining expression profiles of the associative striatum nucleus (6705 annotated genes, p<0.05) and of the hippocampus (over 4000 genes, p<0.05) of SZ patients.

The extended gene lists of annotated genes obtained from the analysis of each dataset (see supplemental file 4) were then used for searching the selected 42 candidates specified in the previous section. Overall, we found significant differential expression values for over 75% (32 out of 42) of the common candidates for SZ, domestication, and/or NC development and function, as discussed above, namely: *BDNF, BMPR1B, CACNA1D*, *CDH2*, *COX4I1, DLGAP1, DLX5, DLX6*, *FOXD3*, *GDNF, GLI3, GRID1, HES1, IMMP2L, JPH3, KDM6B, KIF1B, MSX1, NF1, NIPBL, OLIG2, POMT1*, *RET*, *ROBO2, SKI*, *SOX9*, *SYNJ2, TBX1, TPH1, VDAC1, WNK2* and *ZEB2*. These genes resulted differentially expressed in different brain areas (Figure 4a), the largest number being found within the expression profiles of the frontal cortex (21 genes), the associative striatum nucleus (16 genes), and the hippocampus (11 genes). These functional events are obviously observed in adult (cadaveric) specimens, hence they could not reflect the molecular events that have occurred during early neural development, which are crucial for the etiopathogenesis of SZ. Nonetheless, some insights into the molecular networking that underlie the impaired cognitive and social scenarios acting in the SZ brain could be gathered. Moreover, it is important to note that all these brain areas exhibit anomalies (structural and functional) in schizophrenics, play a role in language processing, and show differences in domesticated animals compared to their wild conspecifics.

**Figure 4.**
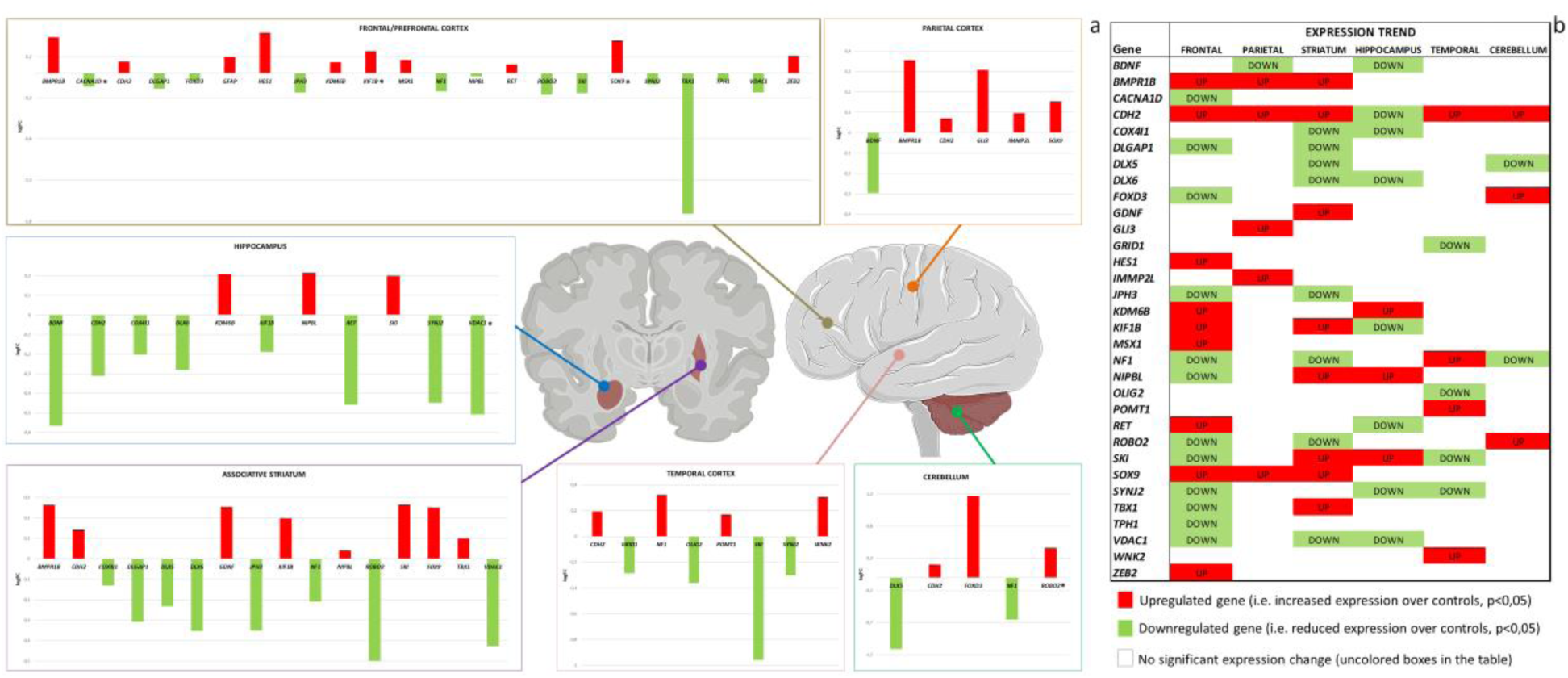
Brain gene expression profiles in SZ patients. The figure on the left (a) illustrates the differential expression of the candidate genes found overlapping between SZ and domestication and/or NC development and function in the tested brain areas. The background diagrams show a coronal section across the midline (on the left) and a lateral view (on the right) of the brain. The pictures were gathered and modified from “Slide kit Servier Medical Art” (available at www.servier.com). In each bar graph: the Y-axis shows the log fold change (logFC) values calculated between SZ patients and controls in each tested dataset (red bar: upregulated gene, green bar: downregulated gene; see text for details). The logFC value of genes labeled with an asterisk was calculated as the mean value of duplicated probeset raw data found in the corresponding dataset (see Supplemental file 4 for extended gene lists). FC values cannot be directly compared when obtained from different datasets (refer to Table 1), as they represent relative expression quantities, normalized over different controls samples in different studies. The expression trends of each gene across the different areas is shown as a cluster view in the image on the right (b).

In the frontal cortex, most of the core-set genes in the gene network displayed in Figure 3 are upregulated, namely *BMPR1B*, *CDH2*, *KDM6B, MSX1*, *RET*, *SOX9* and *ZEB2*. A pivotal role in this selected group is played by *SOX9*, whose activation occurs downstream to BMP signaling (Liu et al., 2013b), consistent with the coherent upregulation *BMPR1B*. KDM6B binds and cooperates with *SOX9*. *SOX9* promotes *CDH2* activation (Sasai et al., 2014), which, in turn, induces the SMAD cascade and leads to *ZEB2* transcriptional activation (Bedi et al., 2014). In the same brain area, *FOXD3* (a *SOX9* repressor, based on String10 prediction) is downregulated, along with other 11 candidates (Figure 4a). Interestingly, genes that displayed differentially expression in the frontal cortex of domestic animals (dogs, pigs, rabbits, and guinea pigs) compared to their wild counterparts included a widely conserved *SOX* gene (Albert et al, 2012), strongly supporting the central role of these gene family in the brain changes implicated in the domestication process.

In the temporal cortex, *OLIG2* and *SKI*, among others, show a reduced expression trend (Figure 4a), possibly indicative of the impaired glial function. Indeed, *SKI* methylation was described in SZ brains (Montano et al., 2016).

It is worth noting that structural and functional anomalies in the frontal and temporal cortices of SZ patients are believed to impact greatly on their language abilities. Accordingly, lexical processing in SZ entails abnormal oscillatory patterns in the left frontal-temporal areas, specifically reduced temporal lobe *α* and left frontal lobe *β* activity (Xu et al., 2013, Sun et al., 2014). Likewise, complex sentences understanding involves reduced activation in the right posterior temporal and left superior frontal cortex in schizophrenic patients (Kircher et al., 2005). Reduced *α* and *β* in left temporal-parietal sites, along with reduced *θ* at right frontal lobe sites is observed during phrase structure chunking (Ferrarelli et al., 2012, Xu et al., 2012). Interestingly too, the ratio fronto-parietal vs. fronto-temporal connectivity has increased from monkeys to apes to modern humans (Hecht et al., 2013), providing the evolutionary scaffolding for our imitation abilities, which underlie cultural innovation (see Boeckx and Benítez-Burraco 2014a for discussion).

Also in the striatum, key candidates, including *BMPR1B*, *CDH2*, *SOX9*, *GDNF* and *SKI* are coherently upregulated, while *DLX* genes and *ROBO2* are downregulated, suggesting that key molecular markers of NC-derived structures are impaired in this nucleus. The striatum is, indeed, part of the cortical-subcortical loop involved in learning speech sequences for articulation, and in extracting complex regularities in auditory sequences (Krishnan et al., 2016).

With regards to the hippocampus, we have found that eight candidates are downregulated, including *BDNF*, *CDH2*, *DLX6*, and *RET* (Figure 4a). Based on the reciprocal functional connections among these genes, discussed above, their shared expression trends may reflect an altered hippocampal synaptic function. Also *VDAC1* and *COX4I1* are downregulated in this brain region (Figure 4a); these genes are both expressed in the mitochondrion and are involved in energy production and ion homeostasis (Richter et al., 2010, Shoshan-Barmatz et al., 2015). Hence, their downregulation reasonably impacts on brain bioenergetics, in SZ and also in other disorders featuring cognitive impairment (Rosa et al., 2016). Interestingly, *KDM6B*, a target of activin signaling involved in cognitive function and affective behavior (Link et al., 2016), is also upregulated in the SZ hippocampus (Figure 4a). The hippocampal expression of SKI gene is increased as well, suggesting aberrant cortical connections in the SZ hippocampus. The hippocampus is involved in language learning (Krishnan et al., 2016), and displays a reduced volume in schizophrenics (Hirayasu et al., 1998), which can be related to phrase chunking difficulties, due to oscillations generated in this region (Murphy, 2015). Interestingly, some domestic animals exhibit changes in the hippocampus that can berelated to differences in their cognitive and behavioral features (Rehkämper et al., 2008).

Finally, few candidates show expression changes in the cerebellum; these include *CDH2*, *FOXD3* and *ROBO2*, coherently upregulated (Figure 4a), confirming their synergic functions, as gathered from the network analysis (see Figure 3). The cerebellum is involved in both motor and non-motor language-related processes, (Mariën et al., 2014, Noroozian 2014), with structural and functional anomalies documented in SZ patients (Keller et al., 2003, Chen et al., 2013)

Taken together, the expression levels of the entire gene set across the tested brain areas suggest that a hallmark of the SZ brain molecular signature is the abnormal activation of the BMP-SOX9-CDH2 axis, given the reproducible upregulation of *BMPR1B*, *CDH2*, *KDM6B*, *SOX9* (in at least two areas; Figure 4b). Conversely, our data show a prevalent repression of *BDNF, DLX5-6, JPH3 COX4I1, DLGAP1, NF1, SYNJ2* and *VDAC1* in the SZ brain, suggesting the overall impairment of molecular mechanisms affecting neuronal survival, synaptic plasticity and functional, and social cognition (Li et al., 2016, Petrella et al., 2016). We reasonably expect that these changes account in part for the specific cognitive, language, and social phenotype of schizophrenics.

Regarding the biological consequences of this overlapping between domestication and/or NC candidates, and SZ candidates, we wish to add two notes of caution. First, we have found that between 20-30% of candidates for domestication/NC are also related to SZ, whereas roughly 5% of the coding genes of the human genome have been implicated in the disease (according to the Schizophrenia Database). Nonetheless, because of pleiotropy, it may well be that some of these common candidates are not playing the same biological function domestication/NC and in SZ. This possibility has to be confirmed experimentally for each candidate. That said, we have shown that some promising functional overlapping can be found. SZ is a late-developing condition, because important brain changes resulting in the disease (e.g. changes in neural pruning mechanisms) occur during adolescence, whereas domestication mostly results from changes in early developmental stages. Nonetheless, available evidence also suggests that the brain and the cognition of SZ patients develop differentially (compared with typically-developing people) since the very beginning, as shown by studies in presymptomatic patients (Cannon et al., 2015; Liu et al., 2015; Filatova et al., 2017; Sugranyes et al., 2017). At the same time, developmental changes, brought about by the early disturbance of the NC function (and by domestication in general), are expected to have an impact throughout lifespan, particularly, because of its effect on the environment. Accordingly, it is reasonable to claim the existence of a biological overlapping between the etiology of SZ, the role of NC in development, and domestication of the human phenotype.

## 5. Schizophrenia and (the evolution of) human language

Many of the genes gathered in our selected gene list play a role in the etiopathogenesis of phenotypes affecting the language domain, reinforcing the link between domestication, SZ, and language. In most cases, though, the implication in neural/cognitive functions could be viewed as a side-effect of the wide pleiotropism and extremely heterogeneity of the vast group of genes considered to lay the polygenic foundation of SZ.

*DLX* genes, including *DLX5* and *DLX6* are also important for the evolution of language-readiness, based on their interaction with *FOXP2* and *RUNX2* (Boeckx and Benítez-Burraco, 2014a). Dlx5/6(+/−) mice show reduced cognitive flexibility that seemingly results from an abnormal pattern of γ rhythms, caused by abnormalities in GABAergic interneurons: this phenotype recapitulates some clinical findings of SZ patients (Cho et al., 2015). During language processing γ rhythms are hypothesised to generate syntactic objects before β holds them in memory and they also contributes to lexical processing (see Murphy 2015 for details). As noted in the previous section, in schizophrenics reduced γ activity is observed at frontal sites during semantic tasks; likewise, higher cross-frequency coupling with occipital α is usually detected (Murphy and Benítez-Burraco, 2016a).

Also *ROBO* genes are core candidates for language evolution (Boeckx and Benítez-Burraco 2014b). *ROBO1* is a candidate for dyslexia and speech-sound disorders (Hannula-Jouppi et al., 2005, Mascheretti et al., 2014), and is involved in the neural establishment of vocal learning abilities (Wang et al., 2015). The *ROBO2* locus is also in linkage with dyslexia and speech-sound disorder and reading (Fisher et al., 2002, Stein et al., 2004), and is associated with expressive vocabulary growth in the unaffected population (St Pourcain et al., 2014).

*GLI3* is involved in a craniofacial syndrome involving cognitive impairment, both in humans (Greig cephalopolisyndactily, OMIM#175700) and in mice (Veistinen et al., 2012; Lattanzi et al, in press; Tabler et al., 2016) with cognitive impairment, which entails language delay (McDonald-McGinn et al., 2010, Lattanzi, 2016). Indeed, it regulates skull development acting on the DLX5/RUNX2 cascade (Tanimoto et al., 2012), hence it is expected to have played a role in the physiological events leading to globularization, in which these genes were seemingly involved (Boeckx and Benítez-Burraco, 2014a; Nearly 98% of Altaic Neanderthals and Denisovans gained a non-synonymous change in GLI3 that is described as mildly disruptive (Castellano et al., 2014).

*OLIG2* confers susceptibility to SZ, alone and as part of a network of genes implicated in oligodendrocyte function (Georgieva et al., 2006). *OLIG2* is also associated with psychotic symptoms in Alzheimer’s disease (Sims et al., 2009) and is up-regulated in the cerebellum of ASD patients (Zeidán-Chuliá et al., 2016).

*SKI* is mutated in Shprintzen-Goldberg syndrome that features skeletal abnormalities and intellectual disability, including speech and language impairment (Au et al., 2014). Speech impairment has been hypothesized to result from poorer phonological abilities, although changes in the quality of the voice (pitch, nasalization) are also observed in patients (Van Lierde et al., 2007).

A well-known polymorphism of *BDNF* (Val66Met) seemingly influence in the pattern of brain activation and task performance during reading, including reading comprehension and phonological memory (Jasińska et al., 2016). This and other *BDNF* polymorphisms have a proven to impact on the language performance of SZ patients (Kebir et al., 2009, Zhang et al., 2016). BDNF is also mentioned as one of the genes defining the genetic architecture of human developmental language impairment (Li and Bartlett 2012).

In Mowat-Wilson syndrome, due to *ZEB2* mutations, severe impairment of productive language is described (Adam et al., 2006, Garavelli et al., 2016).

*NF1* is mutated in neurofibromatosis type 1 (OMIM#162200), a neurogenetic disorder comprising an increased risk for learning and intellectual disabilities, among other major symptoms (Anderson and Gutmann 2015). Affected children may exhibit high rates of social impairment that impact social interaction and skills (Allen et al., 2016; Brei et al., 2014). These might result, in part, from a generalized deficit in the “Theory of Mind” (crucial for language acquisition), which seems to be independent of their general cognitive abilities (Payne et al., 2016). They also feature poor expressive language and preliteracy skills (Lorenzo et al., 2013, 2015). Deficit in fine motor skills usually co-occur, dut to the impairment of fronto-striatal-cerebellar loop (Iannuzzi et al., 2016). Interestingly, heterozygous *Nf1* (+/−) mice show larger brain volumes in the prefrontal cortex, in the caudate and the putamen (part of the language structural network), and in regions involved in social recognition and spatial learning (Petrella et al., 2016). Although these children frequently exhibit ASD symptoms, they outscore IQ-matched children with ASD in eye contact, behavior, and language skills (Garg et al., 2015).

Other genes, not clearly interconnected in the network (see Figure 3), are also clearly implicated in neural functions that are relevant to the language domain.

*NIPBL* is a candidate for Cornelia de Lange syndrome (OMIM#122470), in which the intellectual disability greatly impacts on the expressive language abilities (Boyle et al., 2015, Parisi et al., 2015). The gene is also a putative candidate for childhood apraxia of speech (Peter et al., 2016).

*VDAC1* may reasonably represent a marker of neuronal vulnerability and cognitive impairment, in neurodegenerative conditions including Alzheimer disease (Rosa et al., 2016), and neuronal ceroid lipofuscinoses (Kielar et al 2009). These conditions are characterized by speech and language problems, with a decline in verbal abilities over time (Lamminranta et al., 2001, von Tetzchner et al., 2013).

The region containing *GRID1* has been associated with the parieto-occipital 10-Hz rhythmic activity (Salmela et al., 2016). This α rhythm, which synchronizes distant cortical regions, is involved in lexical decision making and contributes as well to the embedding of γ rhythms generated cross-cortically in order to yield inter-modular set-formation during language processing (see Murphy 2015 for details). This rhythmic activity is found altered in SZ during lexical and sentence processing (Murphy and Benítez-Burraco 2016a).

*IMMP2L* is a candidate for behavioural disorders, including Gilles de la Tourette syndrome (OMIM#137580) (Bertelsen et al., 2014, Gimelli et al., 2014). In this condition multiple motor and vocal tics are described that impact on language processing (Frank 1978).

Abnormal expansion of repeats in the 3' UTR of *JPH3* have been associated with an allelic variant of Huntington disease (OMIM#606438), in which developmental impairment, including neurologic abnormalities and dysarthria, are featured (Seixas et al., 2012, Mariani et al., 2016). Likewise, mutations in *JPH3* give rise to generalized cerebral atrophy, mostly impacting on the basal ganglia (a core component of the brain language circuitry), and are associated with cognitive decline and psychiatric features, including lack of speech due to akinexia (Walker et al., 2003, Schneider et al., 2012).

Also *ABCG1* has been associated with conditions entailing cognitive impairment, language deficits, and neuropsychiatric symptoms (Leoni and Caccia 2015).

*CACNA1D* is associated with several neuropsychiatric conditions (Kabir et al., 2016) and plays important roles in hippocampus-dependent learning and memory (Marschallinger et al., 2015). It also contributes to aversive learning and memory processes (Berger and Bartsch 2014, Liu et al., 2014), and has been linked to neurodegenerative disorders (Berger and Bartsch 2014). Mutations in this gene are considered among risk factors for ASD and intellectual disability, both entailing language deficits (Pinggera et al., 2015).

*HES1* is thought to be important for the evolution of language, because of its specific interactions with *ROBO1* and *RUNX2* (Boeckx and Benítez-Burraco, 2014b).

*POMT1* is associated with clinical conditions entailing severe mental retardation (van Reeuwijk et al 2006, Godfrey et al., 2007, Yang et al., 2016. Mutations in this gene occasionally result in psychotic symptoms (hallucinatory behavior) (Haberlova et al 2014).

*TBX1* and *CRKL* map within the chromosomal hotspot for the Di George/velocardiofacial clinical spectrum, which are neurocristopathies entailing brain anomalies, behavioural disturbances, cognitive impairment, and language delay (Swillen et al., 1997, Swillen et al., 1999, Guris et al., 2001, Glaser et al., 2002). Tbx1 haploinsufficiency in mice causes prepulse inhibition, a robust endophenotype of nonsyndromic SZ (Paylor et al., 2006), and impacts negatively on pup-mother social communication (Takahashi et al., 2016).

Polymorphisms in *NEUROG1* have been associated with SZ and schizoaffective disorder, one of which being significantly associated with increased cerebral gray matter and generalized cognitive deficits (Ho et al., 2008).

*SYNJ2* is an evolutionary conserved gene, and a putative cognitive candidate, whose variants have been found associated with cognitive ageing in selected populations (López et al., 2012).

Finally, mutations in *GFAP* give rise to Alexander disease (OMIM#203450). Specifically, *GFAP* is upregulated in the left posterior superior temporal gyrus (Wernicke’s area) of schizophrenics, a region crucially implicated in language processing (Martins de Souza et al., 2009).

The involvement of most of these common candidates for SZ, domestication and NC in language function provides with an unexpected, intriguing window on language evolution. As we pointed out in the introductory section, an evolutionary link has been claimed to exist between the origins of language and the prevalence of SZ among human populations, because of the nature of the brain changes that brought about language, which plausibly favour the dysfunctions typically found in SZ. In this paper we have argued for putting the focus not only on the split between extinct hominins and AMHs, but also on the time period following the emergence of our species. The main reason is that changes in the social environment linked to our subsequent self-domestication are expected to have contributed to the emergence of modern languages. As we have showed in the paper, many candidates for domestication and NC development and function are involved in language, but also in the etiopathogenesis of SZ, reinforcing the view that domestication, language evolution, and SZ are intimately related. The genes we highlight here might have contributed to this set of late, domestication-related changes in the human phenotype. Interestingly, signals of ancient selection (occurring >1,900 generations ago, prior to the split of present-day human groups) have been found in some of our candidates, particularly, in *ZEB2*, but also in *BMPR2*, a gene encoding another BMP receptor (Zhou et al., 2015). This is in line with the finding that SZ risk alleles may have mainly appeared during this late period, after the emergence of our species. We expect that these recent changes in our candidates contributed as well to the changes that are concomitant to the domestication process.

## 6. Conclusions

Taken together, the data discussed in this paper may provide original hints towards the clarification of some aspect of SZ etiopathogenesis, balancing genetic, epigenetic and environmental factors, and merging development and evolution. The proposed approach may help to disentangle as well the evolutionary history of human cognition, and specifically, of the human faculty of language. In particular, it supports the view that changes in the social context linked to self-domestication contributed decisively to the emergence of modern language and present-day complex languages and that both genetic and environmental factors play a role in this process.

## Acknowledgements

Preparation of this work was supported in part by funds from the Spanish Ministry of Economy and Competitiveness (grant numbers FFI2014-61888-EXP and FFI-2013-43823-P to ABB), and in part by "Linea D1- 2016 and 17" intramural funds from Università Cattolica S. Cuore (to WL).

### Conflicts of interest

The authors declare no conflict of interest.

